# Cardiac contraction velocity has evolved to match heart rate with body size through variation in β-cardiac myosin sequence

**DOI:** 10.1101/680413

**Authors:** Chloe A. Johnson, Jake E. McGreig, Carlos D. Vera, Dan P. Mulvihill, Martin Ridout, Leslie A. Leinwand, Mark N. Wass, Michael A. Geeves

**Affiliations:** School of Biosciences, University of Kent, Canterbury, UK; School of Mathematics, Statistics and Actuarial Science, University of Kent, Canterbury, UK; BioFrontiers Institute and Department of Molecular, Cellular and Developmental Biology, University of Colorado Boulder, Colorado, USA

**Keywords:** evolution, motility, muscle

## Abstract

Heart rate and the maximum velocity of contraction of striated muscle are inversely related to species size. As mammals evolve to different sizes, adaptations are required such as slower contracting heart and skeletal muscles. Analysis of the motor domain of β-myosin from 67 mammals from two clades identifies 14 sites, out of 800, strongly associated with body mass (p<0.01) but not with the clade (p>0.05). Nine of these sites were mutated in the human β-myosin to make it resemble the rat sequence. Biochemical analysis revealed that the rat-human β-myosin chimera functioned like the native rat myosin with a two fold increase in both motility and in the rate of ADP release from the actin.myosin cross-bridge (the step that limits contraction velocity). Both clades use the same small set of amino acids to adjust contraction velocity, suggesting a limited number of ways in which velocity can be manipulated.

---

Proteins adapt and evolve over long time periods tuning their function to the specific needs of the organisms in which they are expressed. Understanding how proteins adapt to different physiology is one of the challenges of current molecular and structural biology. One approach is to consider a protein expressed as different isoforms within a species (paralogues) or in different species (orthologues) where adaptation has taken place. Such a study is easier if a close link can be established between different phenotypes in the organism for which a specific protein function is needed. Striated-muscle myosin motors represent a protein family where such evolutionary relationships can be explored. The maximum contraction velocity of a muscle, V_0_, (along with power output and the velocity at which power and efficiency are maximal) is a key attribute of muscle contraction ^1^;^2^, which is expected to be under selective pressure. The maximum shortening velocity of a muscle is a function of the myosin isoform expressed in the muscle ^3^. In mammals, there are 10 different striated muscle myosins, each expressed from a different gene; many of which have been shown to have distinct biochemical and mechanical properties ^3–6^.

Larger mammals tend to have slower contracting muscles than small mammals where the movement of a larger mass results in slower velocity. This phenomenon is well-established in heart muscle where heart rate is inversely related to body mass for a wide range of species ^7^. If velocity of contraction is matched to the size of the animal expressing the myosin, then there should be changes in the myosin sequence to tune the myosin properties to the species size and correspondingly, to the physiology of the muscle. In a recent study we tested the hypothesis that muscle myosin-II isoforms from mammals would have adaptations in protein sequence associated with mean body mass ^8^. In this study, of ∼730 sequences from 12 myosin-II isoforms from an average of 65 mammalian species, there was a strong correlation of the number of sequence changes with differences in species mass. The correlation was strongest in adult striated muscle myosins IIa and IIb and β. β-myosin is found in the heart and in slow Type I muscle fibers. Here we examine in greater detail the sequence differences within the mammalian β-myosin to establish the relationship between sequence and velocity of contraction.

The contractile properties of mammalian muscles have been widely studied but detailed mechanical and biochemical studies have been completed on only a few species and myosin isoforms. Such data are available for Type 1/β-cardiac/slow muscle fibres from four species (see Fig 1). Each of these muscles expresses only the β-myosin isoform (MYH7), and each species has a characteristic contraction velocity that varies approximately five-fold across the set of four muscles. Moreover, it is well established for the β-myosin isoform that the contraction velocity is limited by how fast ADP escapes from the actin.myosin cross-bridge after the working stroke is completed ^9^. Data in Fig 1 show that like velocity, the ADP release rate constant differs 4-5 fold across the set of four myosins. Furthermore, the measured rate constant for ADP release from the actin.myosin complex, measured using the purified β-myosin isoform, is exactly that predicted to limit the contraction velocity (based on the detachment limited model of contraction (see Fig 1 legend) ^9–11^. It is therefore expected that this set of myosins will have changes in the amino acid sequence that alter the ADP release rate and hence contraction velocity.

**Figure 1.**
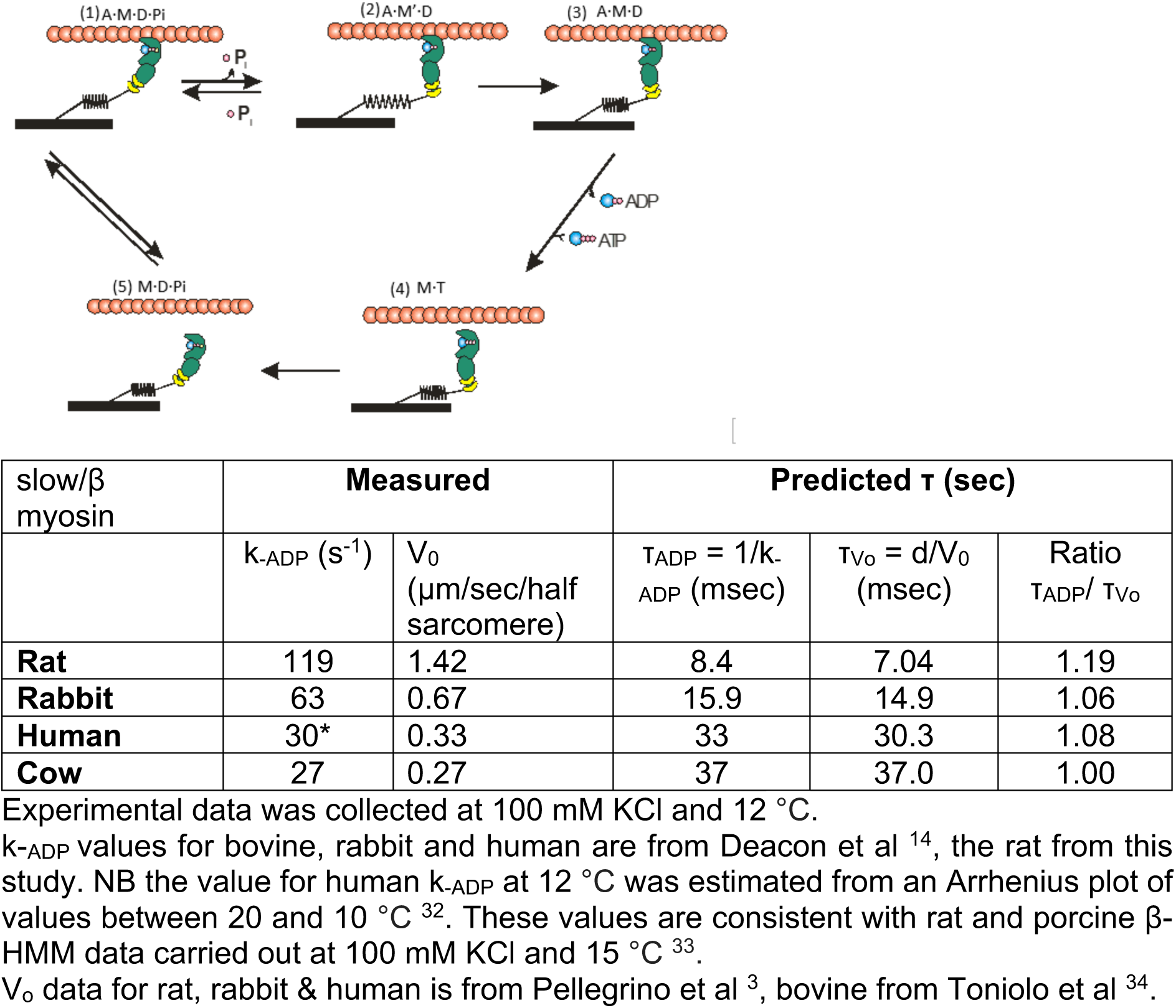
The relationship between the predicted and measured parameters for four slow/beta cardiac myosin isoforms. Figure adapted from ^30^. In terms of the actin myosin cross bridge cycle the dominant model proposes that the maximum velocity (V_0_) is limited by the lifetime of the strongly attached force holding state (τ) the “detachment limited model” (V_0_ = d/τ where d is the working stroke of the cross bridge; assumed here to be 5 nm ^10^. For the mammalian, β-cardiac/slow-muscle myosin isoform, it is well established that τ is defined by the rate constant controlling ADP release k_-ADP_ = 1/τ (see table below). Thus, values of k_-ADP_ measured using myosin motor domains isolated from β-cardiac/slow muscle of a mammal, predict remarkably accurately the maximum shortening velocity of a muscle fibre taken from the same tissues.

Examining the sequences of the 800 amino acid motor domains of the four β-myosins in Fig 1 show them to be 96% identical which means there are 49 sequence differences among the four, with 34 differences between rat and human. These differences in sequence are scattered throughout the motor domain (see Fig 2 and alignment in Fig S1). It seems likely that groups of amino acids and not a single amino acid change determine the functional differences between the β-myosins. If the correlation between body size and contraction velocity is a general phenomenon amongst mammals, as is seen for resting heart rates ^7^, then a bioinformatics study of β-myosin sequences would be a way to identify which sequence changes correlate with size. We hypothesised that the variation of β-myosin contraction velocity (and rate of ADP release from the cross bridge) with the size of the mammal is due to a subset of the sequence changes observed for different mammals, and that these tune myosin velocity to that appropriate for the size of species. Here, we examine a set of 67 mammalian β-myosin sequences ^8^ using a bioinformatics approach to identify a group of 12 amino acids which have the strongest association with the size of the mammal. Of these 12 amino acids, nine differ between human and rat β-myosin and we test our hypothesis through the construction and subsequent biochemical characterisation of a rat-human β-myosin chimera (hereafter referred to as chimera).

**Figure 2.**
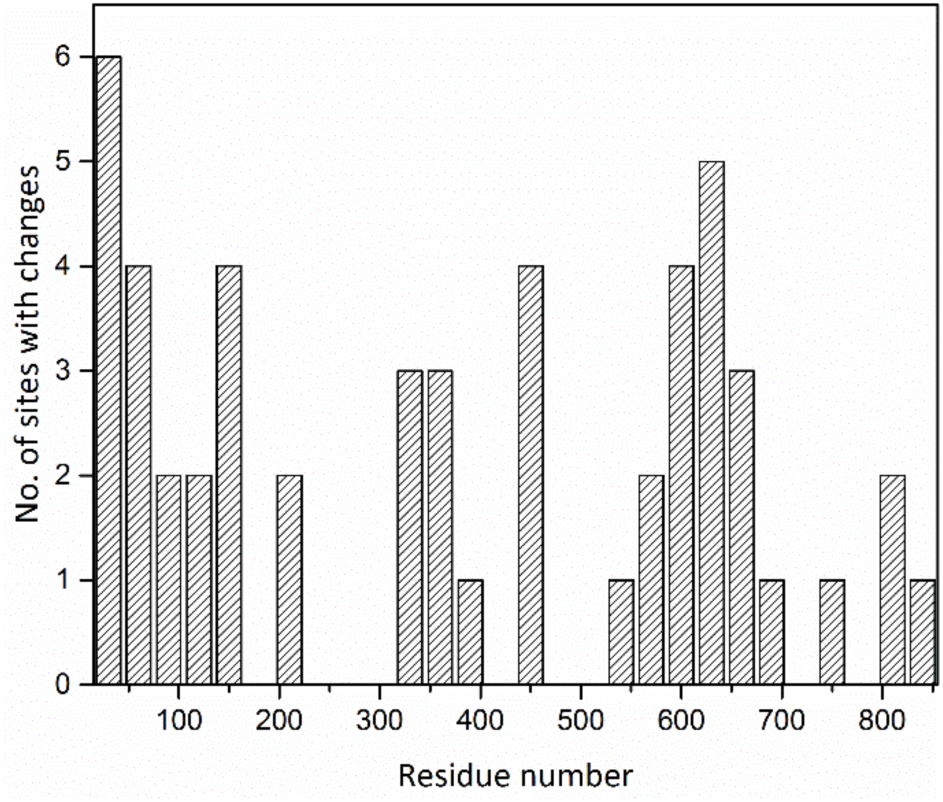
Distribution of sequence variations for the β-myosin sequences listed in Figure 1. The sequence which is ∼96% conserved was divided into blocks of 30 residues and the no of sites within that group showing a change is plotted. The maximum number is six residues in region 1-30 reflecting the high degree of identity among the four sequences.

## Methods

### Sequence Analyses

Amino acid sequences for the β-myosin motor from 67 different mammalian species were aligned using Clustal Omega ^12^ comprising organisms from the clades Euarchontoglires (32), Laurasiatheria (30), Metatheria (4) and Afrotheria (1). The start and end points of the motors in *Homo sapiens* were considered to be residues 1 and 800 (based on UniProtKB – P12883).

### Statistical analysis

For each position in the alignment that had more than one amino acid present, species masses were compared between the two highest frequency amino acids at that position using the Mann-Whitney U test, a nonparametric two-sample test. Multiple testing was accounted for by applying the Bonferroni correction. To avoid very imbalanced comparisons, the analysis was not run if the frequency of the second amino acid was less than 10% of the frequency of the most frequent one. Where more than two amino acids were present at an alignment position, only the two most frequent amino acids were considered. See Fig S3 for details of sites with more than 2 amino acids.

Alignment positions were divided into three groups: those with an adjusted p-value (p_adj_) less than 0.01 (with Bonferroni correction applied this is equivalent to a p-value of p= 9.50×10^-04^), those with 0.01 < p_adj_ < 0.05 (5% significance threshold equivalent to p=9.62×10^-4^), and those with a p_adj_ >0.05. In addition, the two highest frequency amino acids were coded as 0 and 1 and a logistic regression model was fitted with log(mass) as the explanatory variable (Fig 3 & Supplementary Fig 2). In order to overlay these residue plots, as the coding of the amino acids as 0 and 1 was arbitrary, it was done in such a way that the slope of the fitted logistic regression line was positive (Fig 3). The value of mass at which the two amino acids were predicted to be equally likely to occur was estimated from the regression line.

**Figure 3.**
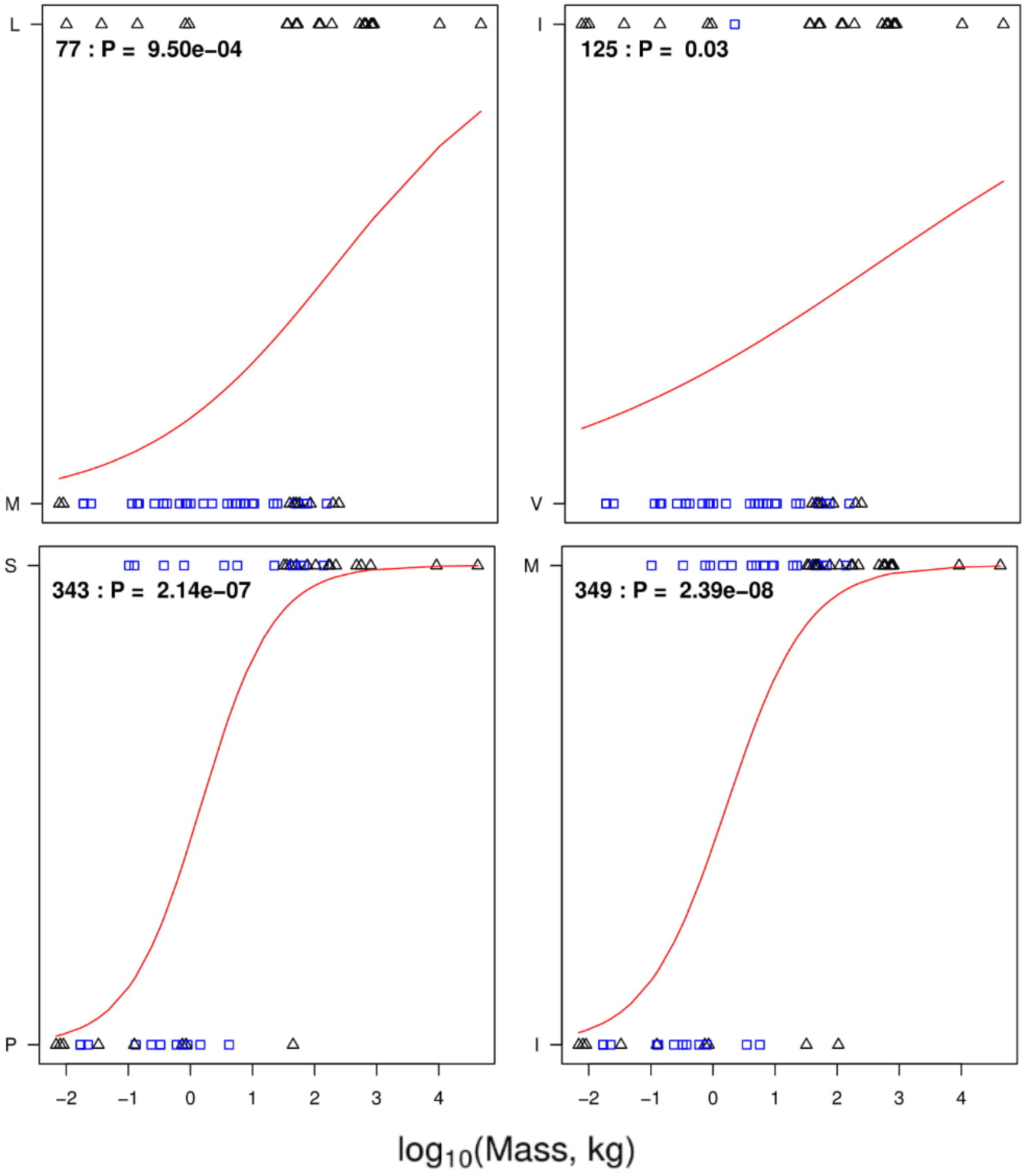
Residue-mass transition plots. Binomial regression mapping the transition of the most frequent amino acid at positions in the motor region of β-myosin to the second most frequent amino acid at that position. The residue numbering is that of the human β-cardiac myosin, as oppose to the alignment position. The blue squares are Euarchontoglires, and the triangles are Laurasiatheria. The P-value with each plot arises from a test of the null hypothesis that amino acid type is unrelated to mass.

For each alignment position, a 2×2 table was constructed classifying the species by amino acids present (most frequent and second most frequent) and clade. Fisher’s exact test was used to test whether these two factors were associated. The residue and -log_10_ of the P-value from the Fisher’s exact test were plotted to identify residues for which the amino acid variation was likely to have resulted from clade associated changes. The residue and -log_10_ of the p-value from the Mann-Whitney U test were also plotted to determine when residue variation was likely attributed to mass changes. Finally, the -log_10_ p-values obtained from both tests were plotted against each other. For each of these plots, lines at positions of the Bonferroni adjusted p-values 0.01, and 0.05 were added to assess the confidence in each residues association with mass or clade.

All statistical analyses were run in R ^13^.

### Molecular Biology of the chimera

A pUC19 plasmid containing the human β-myosin motor domain gene was digested with NsiI and NgoMIV to excise DNA encoding for residues 310 – 599 of the human β-myosin. This region was replaced with a complementary pair of synthetic oligos encoding for the same region, but with the nine amino-acid substitutions listed (Ala326Ser, Ser343Pro, Leu366Gln, Ile421Ala, Thr424Ile, Ala430Ser, Arg434Lys, Phe553Tyr, Pro573Gln). The subsequent clone was confirmed by sequencing. This chimera gene was cloned into a pShuttle CMV vector to allow recombinant replication-deficient adenovirus production, as previously described (Deacon et al 2012).

### Protein purification

The chimera and the human β-myosin motor domains (known as subfragment 1 or S1) were expressed and purified as descried previously ^14^. Briefly, the adenoviruses were used to infect C_2_C_12_ myotubes in culture and resulted in overexpression of recombinant myosin proteins. The heavy chains (residues 1-842) were co-expressed in C_2_C_12_ myotubes with His-tagged human ventricular essential light chain. The recombinant proteins also carried the endogenous mouse regulatory light chain (MLC3). This is homologous to subfragment 1, S1, generated by proteolytic digestion of myosin. For motility assays the heavy chain was additionally tagged with an eight residue (RGSIDTWV) C-terminal extension. Cell pellets were homogenized in a low salt buffer and centrifuged, and the supernatants were purified by affinity chromatography using a HisTrap HP 1 ml column. The proteins were then dialyzed into the low salt experimental buffer (25 mM KCl, 20 mM MOPS, 5 mM MgCl_2_, 1mM DTT, pH 7.0).

The SNAP-PDZ18 affinity tag used for *in vitro* motility measurements were purified as described in ^15,16^. SNAP-PDZ18 was expressed through a pHFT2 expression vector, and the plasmid transformed into *E. coli* BL21 DE3 cells. The protein was purified using nickel-affinity chromatography, and dialyzed in PBS.

Actin was prepared from rabbit muscle as described by ^17^. The actin was labelled with pyrene at Cys-374 as described in ^18^. When used at sub-micromolar concentrations the actin was stabilized by incubation in a 1:1 mixture with phalloidin.

Rat β-myosin S1 was prepared from soleus muscle which was dissected immediately post mortem and stored on ice. The muscle was homogenised into Guba-Straub buffer and left to stir for 30 minutes. After centrifugation at 4600 RPM for 30 minutes at 4 °C, the supernatant was subject to myosin precipitation as described in ^19^. The resulting myosin was digested with 0.1 mg chymotrypsin per ml of solution and left to stir for 10 mins exactly, at room temperature. To stop the digestion, 0.5 mM phenylmethylsulfonyl fluoride (PMSF) was added and the solution left to stir for 10 minutes. The digested myosin solution was dialysed into the low salt experimental buffer overnight (25 mM KCl, 20 mM MOPS, 5 mM MgCl_2_, 1mM DTT, pH 7.0). Precipitated myosin and light meromyosin was pelleted and removed via centrifugation at 12,000 RPM for 10 minutes, with the supernatant containing the purified soleus S1. SDS-Gels of the purified protein were run and compared to the expressed human β-myosin and chimera S1.

### Stopped flow

Kinetic measurements for S1 of chimera, human β-myosin and rat soleus myosin were performed as described previously ^5,14,20^. Solutions were buffered with 25 mM KCl, 20 mM MOPS, 5 mM MgCl_2_, 1 mM DTT at pH 7.0, and measurements were conducted at 20 °C on a High-Tech Scientific SF-61 DX2 stopped-flow system. Traces were analysed in Kinetic Studio (TgK Scientific) and Origin.

### In vitro motility assay

Motility assays were performed essentially as described previously ^16,21^. Briefly, flow chambers were constructed with coverslips coated with nitrocellulose mounted on glass slides. Reagents were loaded in the following order: 1) SNAP-PDZ18 affinity tag; 2) BSA to block the surface from non-specific binding; 3) S1 of human β-myosin or the chimera with an eight amino acid C-terminal affinity clamp; 4) BSA to wash the chamber; 5) rhodamine-phalloidin-labelled rabbit actin; 6) an oxygen-scavenging system consisting of 5 mg/ml glucose, 0.1 mg/ml glucose oxydase and 0.02 mg/ml catalase 7; 2 mM ATP. Partially inactivated myosin heads in S1 preparations were removed by incubating with a 10-fold molar excess of actin and 2 mM ATP for 15 minutes, then sedimentation at 100,000 RPM for 15 minutes. Supernatant was collected containing active myosin heads. All solutions were diluted into 25 mM imidazole, 25 mM KCl, 4 mM MgCl_2_, 1 mM EGTA, 1 mM DTT, pH 7.5. Actin filaments were detected using a widefield fluorescence imaging system (described in ^22^) with UAPON 100XOTIRF NA lens (Olympus) and QuantEM emCCD camera (Photometrics). The system was controlled and data analysed using Metamorph software (Molecular Devices, Sunnyvale, USA). Assays were performed at 20 °C and were repeated with three fresh protein preparations, with at least three movies of 30 second duration, recorded at a rate of 0.46 sec per frame. Individual velocities were determined from motile filaments that demonstrated a smooth consistent movement over 10 frames (4.6 sec). 100 individual measured velocities were used to calculate the mean velocity for each recombinant myosin.

## Results

The alignment of the β-myosin sequences from four mammals (Fig 2 & Fig S1) demonstrates that while the sequences are highly conserved, the 49 sites of variation among the species are scattered throughout the motor domain. High levels of variation are found in the N-terminal domain (1-60) and near the surface loops, Loop 1 (near residue 210) and Loop 2 (near 630). These loops are known to be hypervariable across the larger myosin family ^23^. The broad distribution of the sequence variants means that an experimental approach to define which residue changes are linked to the change in ADP release (and hence velocity of contraction) is too complex to consider. Instead, we used a bioinformatics-based approach to identify the residues most likely to be linked to the change in velocity of contraction.

### Distinguishing between variation due to clade and body mass

We analysed 67 complete sequences of the β-myosin motor domains from species ranging in size from 7g (Brandt’s bat, *Myotis brandtiibat*) to 42,000 kg (sperm whale, *Physeter catodon*). Of these, about half were Euarchontoglires (32, e.g. rodents and primates) and half were Laurasiatheria (30, e.g. bats, ungulates, cetaceans). We used this set of sequences to distinguish between sequence changes that had a high probability of association with the clade vs those that correlated with the size of the animal (see Methods). A total of 171 sites were identified where a sequence change occurred. At the majority of these positions, variation occurred in a small number of species and is unlikely to be associated with changes in function relevant to body mass, so 119 positions where a sequence change was present in less than 10% of the species were excluded. This included 84 sites where a change occurred in only one species, while changes in two species occurred at 20 sites and in three species at four sites.

The remaining 52 sites of variation were analysed to distinguish between changes that correlated with clade and those that correlated with body mass (as illustrated in Fig 3 for four sites with the remaining plots in Supp Info – Fig S2). In most cases, only two residues were observed at each specific site; in the small number of cases (11/52) with multiple amino acids, only the two most frequent residues were considered. The identity of the two most frequent amino acids were coded as 0 and 1 and a logistic regression model was fitted with log(mass) as the explanatory variable (Fig 3, Fig S2; see methods) to model the transition between residues. Data for four sites are presented in Fig 3 and of the sites shown, two had a strong correlation with species body mass (the amino acid common in small mammals is given first P343S, I349P; p_adj_ ≤ 0.01. Note the adjusted one percent significance threshold is p= 9.50×10^-4^ and the 5% significance threshold is p=9.62×10^-4^. p_adj_ will be used to indicate the adjusted significance threshold), and each has a distinct midpoint mass for the transition between the two amino acids. In contrast, I125V has a low association with mass (p = 0.03) and M77L has an intermediate association (p= 9.50e-04), however both M77L and V125I have a strong association with the clade (Fig 4); L77 and I125 are found almost exclusively in *Laurasiatheria*.

**Figure 4.**
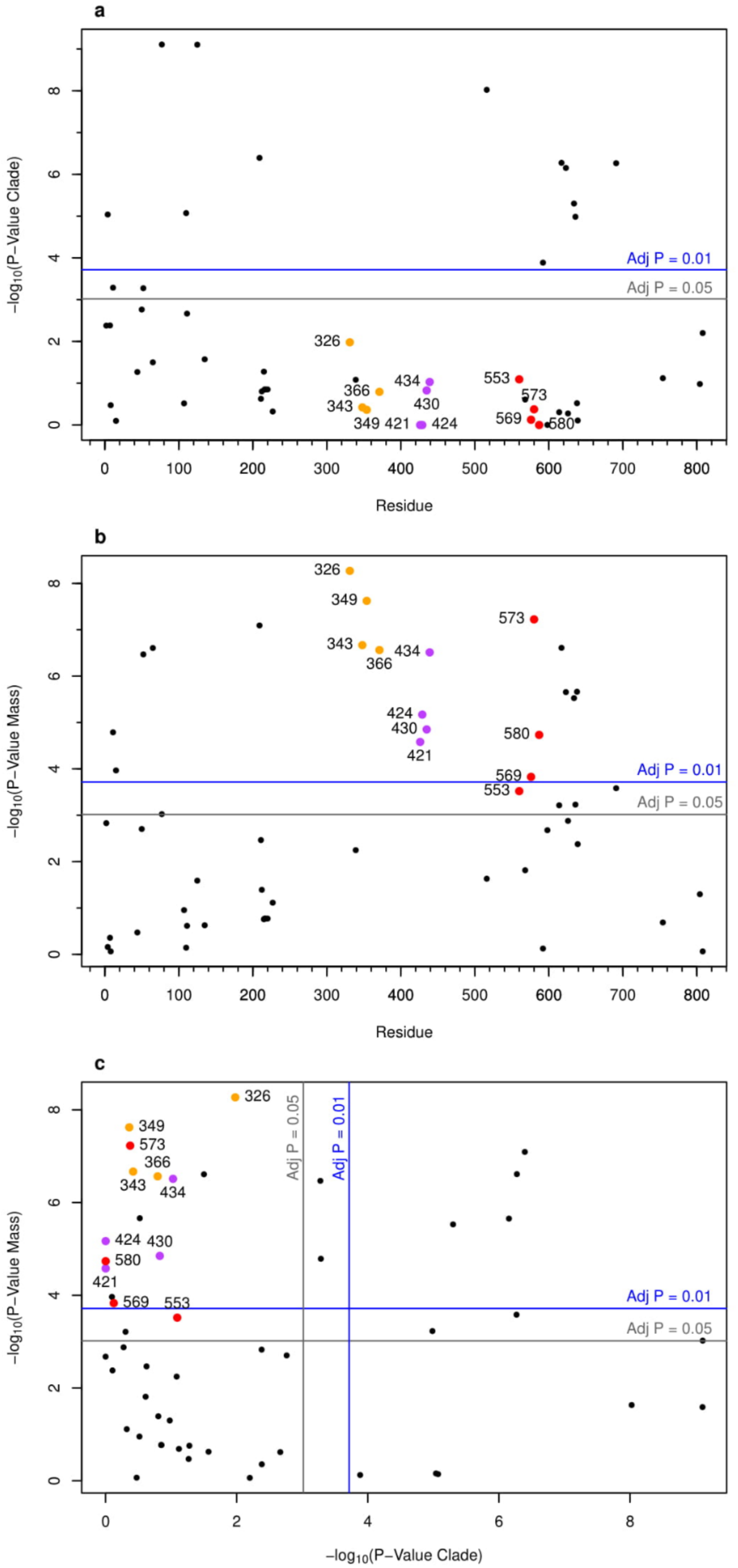
Residue mass change association with mass, clade, and each other. The association of each residue with clade (A), mass (B), and the association of the P-values of clade vs mass (C) are plotted. The significance of the P-values is shown with the Bonferroni adjusted lines drawn onto the plots (blue and grey). The three groups of residues investigated are highlighted in orange, purple and red, and labelled with the human residue numbering in each plot.

Overall, only 12 sites had a very strong association with clade (p_adj_ ≤ 0.01) and a further two were significant at the five percent level (p_adj_ ≤ 0.05; Fig 4A). Some of these residues occur in two groups; one group of four in the N terminal region below residue 135 (4, 11, 52, 77, 110, 125) and four residues near surface loop 2 (610, 616, 627, 629). The remaining four are at D208E, E509T, T585I and I684M. Twenty positions were strongly associated (p_adj_ ≤ 0.01) with body mass and a further four were significantly associated at p_adj_ ≤ 0.05 (Fig 4B). Nine positions were associated with both clade and body mass (Fig 3C), which is likely to represent that the very largest mammals (body mass > 500 kg) in the data set are all *Laurasiatheria* (Fig S5).

Twenty residues had a very strong association with mass, p ≤ 1.92×10^-4^ (1% significance threshold, adjusted for multiple testing) and a further four significant at the five percent level (p < 9.62×10^-4^; Fig 4B). Twelve of these 24 sites occur in the known hypervariable regions, four in the N terminal region (**11**,15,**52**,65, bold **residues** also occur in the clade list), one in loop 1 (**D208E**), and a further six occur in or near loop 2 (607,**610**,**616**,**627**,**629**, 631)) and one at **I684M**. The remaining 12 sites (Table 2) group into three sets of four; most with p_adj_ ≤ 0.01 (coloured in Fig 4 & 5). Comparing the strength of association between clade and body mass, these 12 sites, are strongly associated with body mass but not with clade (Fig 4C). Hence, we propose that these 12 positions are likely to be important in determining the β-myosin velocity of contraction. At eight of the twelve positions only two amino acids are observed, one position contains three amino acids, although the third is only present once (residue 366). Multiple amino acids (4-7) were observed at the remaining three positions. For two of the positions, 421 and 424, this reflects a subset of the species from one clade having an alternate amino acid in some of the larger species (see Sup Fig S3).

The first group of residues is in a region 331-371 (Orange in Fig 4 and 5) adjacent to the exon 7 region of Drosophila myosin II (and the “linker region”). This region is one of four exons in the single myosin II gene of Drosophila which are alternately spliced to generate all isoforms of myosin II in Drosophila ^24^. We have previously shown ^25^ that the alternatively spliced forms of this region alter ADP release in the Drosophila myosin. The second set (426-439; Magenta in Fig 4 and Fig 5) is in a long helix (Helix-O) in the upper 50 kDa domain that links an actin binding site (the myopathy loop) to the nucleotide binding pocket (via switch 2). The third region (560 - 587; Red in Fig 4 and Fig 5) is in the lower 50 kDa domain and lies close to loop 3, an actin binding site in the lower 50kDa domain.

**Figure 5.**
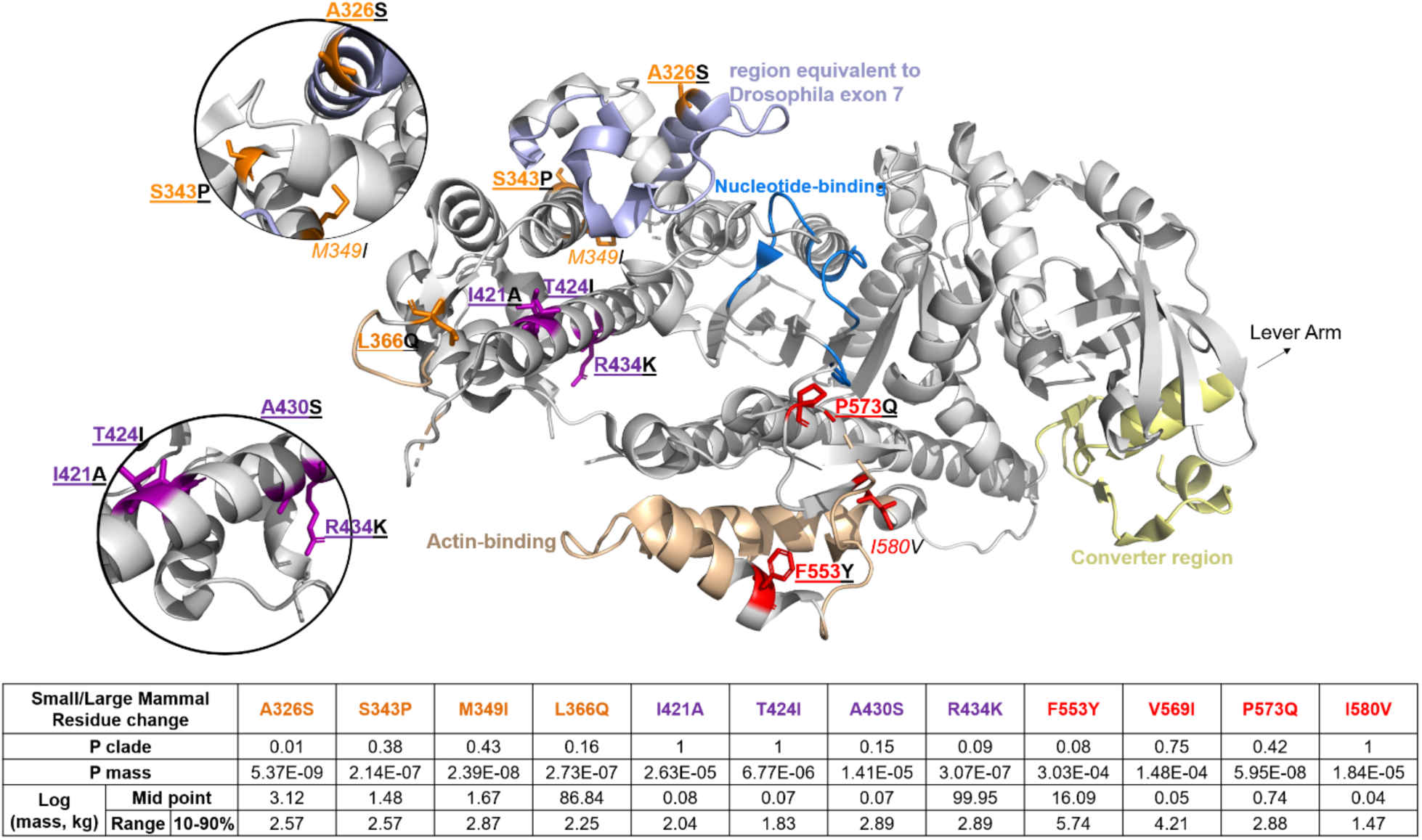
Location of residues switched in the chimera. Structure of human β-myosin (PDB:ID 4DB1). The actin-binding site is highlighted in brown, exon 7 in blue, the nucleotide binding site in marine blue, and the converter region in **yellow**. The three groups of residues investigated are highlighted and labelled in orange (326, 343, 349, 366), purple (421, 424, 430, 434), and red (553, 569, 573, 580) in each plot, and those that were switched are in bold and underlined.

### Experimental testing of the computational predictions

We have previously expressed the motor domain of human β-myosin in mouse C_2_C_12_ muscle cells and isolated the protein using His tags attached to the co-expressed human light chain. This is currently the only way to express mammalian striated muscle myosin motors but is complex and time consuming and yields just a few mg of protein ^6,26,27^. To test the hypothesis that the highlighted group of 12 residues are responsible for a significant part of the adjustment of ADP release rate constant, we generated a chimeric human-rat β-myosin motor domain where the nine positions (of the 12) that vary between human and rat were replaced with the amino acid present in rat (A326S, S343P, L366Q, I421A, T424I, A430S, R434K, F553Y, P573Q – human residue number and amino acid listed first). The other three positions are the same in rat and human (354, 576 & 587). At residue 421 we replaced Ile with Ala as present in rat, although Ser is present in most of the smallest mammals (See Fig S3).

As shown in Fig 1, the velocity of contraction for β cardiac/Type I slow muscle fibres in rat and humans differ by a factor of ∼4. Given that these residues have a range of transition masses (see Fig 6) the hypothesis is that each of these nine residues will contribute a fraction of the difference between the rat and human β-myosin ADP release rate constant and hence, velocity. With all nine residues changed, our prediction is that the differences in the rate constant should be large enough to be easily detectable.

**Figure 6.**
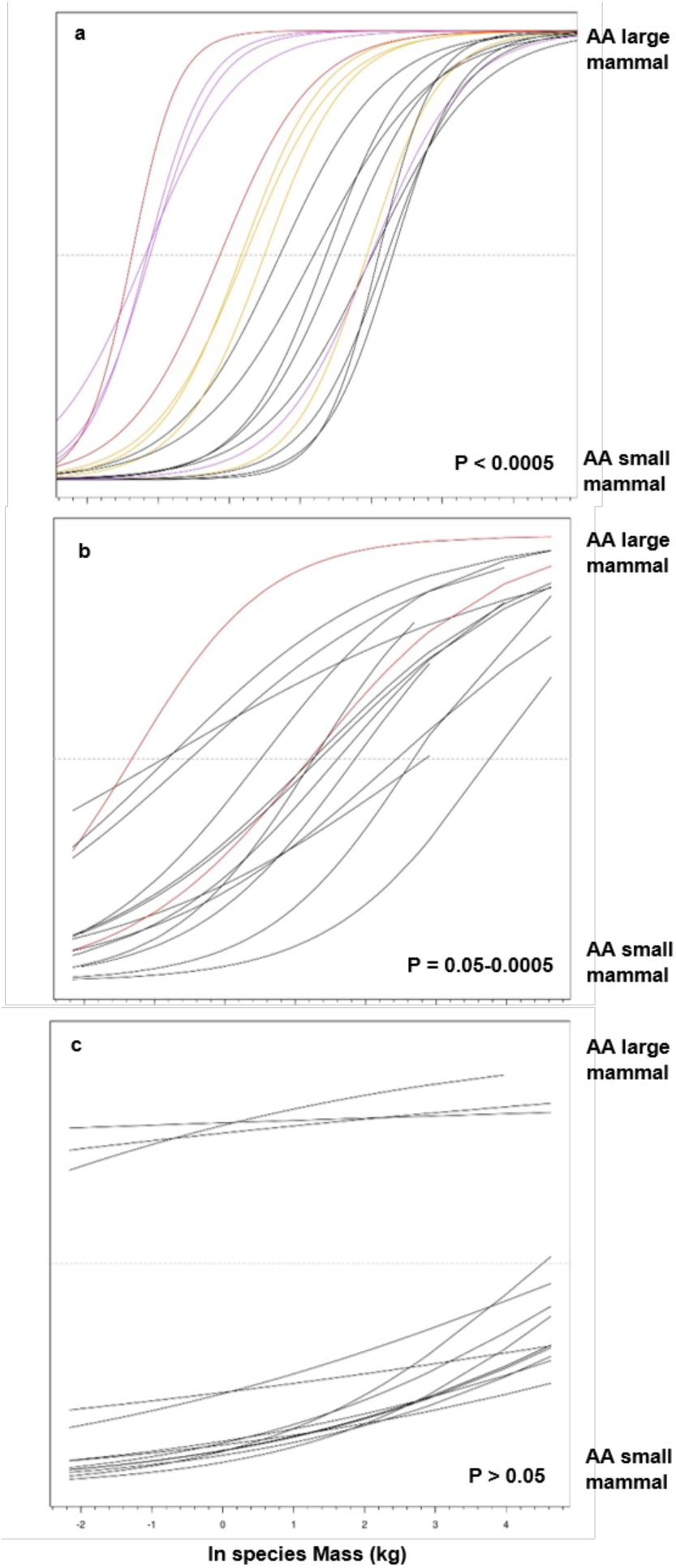
Residue mass transition plots. Overlapping binomial regression mapping the transition of the most frequent amino acid at positions in the motor region of β-myosin to the second most frequent amino acid at that position for residues at three Bonferroni adjusted P-value significance values. These are P < 0.0005 (A), P = 0.05-0.0005 (B), P > 0.05 (C). The amino acids (AA) that occur most frequently and second most frequently are at the extremes of the y-axis. The three groups of residues investigated are highlighted and labelled in orange (326, 343, 349, 366), purple (421, 424, 430, 434), and red (553, 569, 573, 580) in each plot.

The S1 fragment of human β-myosin and the chimera were expressed in C_2_C_12_ cells and purified with the human essential light chain attached (see Fig 5). Few details of the kinetic characterisation of the rat β-myosin S1 have been published ^28^. The rat β-myosin S1 was therefore purified from rat soleus muscle to use as a comparator for the chimera. The supplementary data include the SDS PAGE of all three proteins used in this study and demonstrates that all three proteins are pure and contain the appropriate light chains (Fig S4).

As a test of the behaviour of the chimeric protein, the ATP-induced dissociation of the chimera from pyrene labelled actin was monitored and compared to the recombinant human and the native rat S1. A typical transient is presented inset in Fig 7B and the observed amplitude of the signal change was the same for all three proteins. The similarity of observed amplitudes of the pyrene signal changes for the chimera, human and native rat proteins indicates that the chimera binds actin and releases it on ATP binding as for the human and rat S1. This is consistent with the chimera being a fully folded and active protein. A plot of the observed rate constant (k_obs_) vs [ATP] gives a straight line which defines the apparent 2^nd^ order rate constant for the reaction (Fig 7B) and appears the same for all three proteins. The observed rate constant of this reaction has been defined for many myosins and has two components, k_obs_ = [ATP] K’_1_k’_+2_. The reaction is sensitive to both the affinity of ATP for the complex (K’_1_) and the efficiency with which ATP induced a major conformational change in the myosin (k’_+2_). This involves the closure of switches 1 and 2 onto the ATP and the opening of the major cleft in the actin binding site of myosin. The absence of any change in K’_1_k’_+2_ is consistent with a well preserved nucleotide pocket and a preserved communication pathway between the ATP binding pocket and the actin binding site.

**Figure 7.**
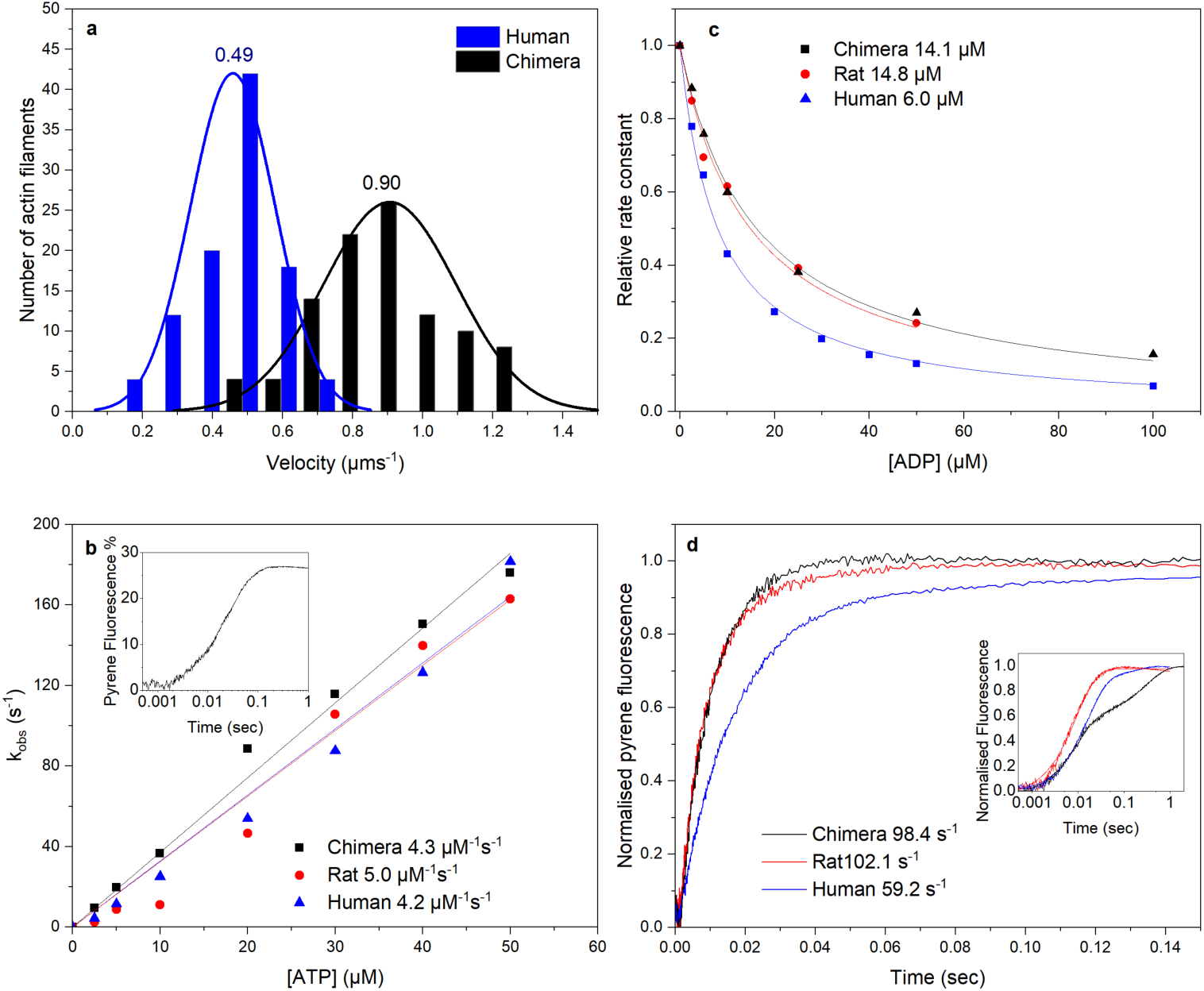
Stopped-flow analysis of the chimera, rat and human β-S1 proteins. A. Histogram of *in vitro* velocity of 100 rhodamine-labelled-phalloidin actin filaments moving over human β-S1 or chimera S1. The solid line shows the data fitted to a single Gaussian curve. The mean velocity for the human β-S1 was 0.49 ± 0.028 μms^-1^ and for the chimera 0.90 ± 0.015 μms^-1^. B. The effect of ATP concentration on k_obs_ for ATP-induced dissociation of pyrene-actin.S1. The gradient generates a second order rate constant of ATP binding; the values for the 3 proteins are highlighted next to the plot. Inset shows an example transient of 50 nM pyrene actin-chimera S1 mixed with 20 µM ATP, resulting in a fluorescence change of 26 %. C. Plot of k_obs_ dependence on [ADP] for the ATP induced dissociation of pyrene-actin.S1. 50 nM pyrene-actin.S1 was mixed with 10 µM ATP and varying [ADP] (0-100 µM). Numbers indicate the values of ADP affinity for acto.S1, k_ADP_, for the 3 proteins. D. To measure k_+ADP_, ADP is displaced from pyrene-actin.S1.ADP complex by an excess of ATP. 2 mM ATP was mixed with 250 nM S1 which was pre-incubated with 500 nM pyrene-actin and 100 µM ADP. k_+ADP_ values for the 3 proteins are given 7D. Inset showing data on a longer log time scale showing the slow phase components of the transients. The average values from 3 independent measurements for experiments shown in B, C and D are summarised in Table 1.

The affinity of ADP for actin.S1 was measured in a competition assay with ATP (Fig 7C) and the affinity of ADP for the rat actin.S1 complex (14 μM) was 2.3 fold weaker than for the human WT protein (6.3 μM). These values are consistent with published values ^29^. The chimera was distinct from the human S1 and indistinguishable from the rat S1. To confirm this result the ADP release rate constant was measured directly by displacing ADP from actin.S1.ADP through addition of an excess of ATP. The results (Fig 7D) for human and rat S1 are again consistent with published values with ADP leaving the rat complex at ∼ 2X the rate of the human complex (107 vs 59 s-1). The chimera was indistinguishable from the rat S1. As predicted, the amino acids introduced into the human β-myosin motor domain weaken ADP affinity for actin.myosin by accelerating ADP release to make the human β-myosin S1 behave like the rat β-myosin S1.

**Table 1.** Comparison of ATP and ADP binding parameters of native rat S1, and the C_2_C_12_ cell expressed human β-myosin and chimera S1. Errors reported are SEM, except for motility which is the HWHM of the normal distribution.

|  | **Rat**<br>**native**<br>**S1** | **Chimera S1** | **Human S1** | **Chimera/Rat**<br>**ratio** | **Chimera/Human**<br>**ratio** |
| --- | --- | --- | --- | --- | --- |
| **ATP binding**<br>**to A.S1**<br>**($\mu\text{M}^{-1}\text{s}^{-1}$)** | 5 ± 0.16 | 4.5 ± 0.2 | 4.4 ± 0.3 | 0.9 | 1.02 |
| **ADP affinity**<br>**to A.S1**<br>**($\mu\text{M}$)** | 14 ± 0.09 | 14 ± 0.14 | 6.1 ± 0.7 | 1 | 2.3 |
| **ADP release**<br>**from**<br>**A.S1.ADP**<br>**($\text{s}^{-1}$)** | 107.2 ±<br>7.3 | 100.7 ± 4.2 | 59 ± 3.3 | 0.94 | 1.71 |
| **Motility**<br>**($\mu\text{m.s}^{-1}$)** | NA | 0.90 ± 0.221 | 0.49 ±<br>0.129 | NA | 1.84 |

Data are from three separate preparation of protein from either different cell pellets (expressed protein) or different soleus muscles from the rat. Experimental conditions were 25 mM KCl, 20°C.

#### Footnote

The inset shown in Fig 7D; a complication of the ADP displacement measurement is that ADP displacement from human β-myosin occurs in two phases (fast and slow). The fast phase corresponds with ADP released at the end of the normal ATPase cycle while the slow phase is a trapped ADP which is released much slower and at a much rate slower than the overall cycling. This is therefore a dead-end side branch of the pathway commonly seen in slow muscle & non muscle myosins ^14,28,30^. The fraction of ADP trapped in this way is characteristic of each myosin. The rat β-myosin S1 has no apparent slow phase, the human has ∼10% of ADP released in the slow phase while the chimera has a larger fraction (∼40%) of the total ADP released in the slow phase. The role of the substituted amino acids in the slow phase requires further study, but the reader is referred to the literature for a broader study of this phenomena.

Motor activity of the recombinant human β-myosin S1 and the chimera protein was measured using an *in vitro* motility assay (Fig 7A). This assay determines the myosin-mediated velocities of fluorescent actin-filaments moving on a nitrocellulose-coated slide surface. The human WT β-myosin moved actin at a velocity of 0.49 μm.s^-1^ at 20 °C.. Introduction of the nine rat amino acids into the WT protein increased the mean filament velocity by almost 2-fold, from 0.49 μm.s^-1^ to 0.9 μm.s^-1^ for the chimera, which is consistent with predictions from the ADP release-rate data (predicted velocity assuming a 10 nm step size 0.59 and 1.00 μm.s^-1^). Our human S1 velocity was similar to the 0.612 μm.s^-1^ value reported by Ujfalusi et al, which was measured at 23 °C ^31^. A similar velocity of 0.378 μm.s^-1^ was reported for full length human-β myosin at 25 °C ^3^. who aslo reported a velocity of 0.624 μm.s^-1^ for the rat. This gives a rat/human ratio of 1.65, very similar to our chimera/human ratio (1.84).. The motility assay was not performed for the rat S1 as we do not have an expression system for the protein. The native rat S1 has only a single light chain and lacks a tag to attach the protein to the surface. The rat protein will not therefore give a valid comparable measurement. However, it is known from the literature (Table in Fig 1 & references therein) that the rat protein moves 3-5 times faster than the human protein, depending upon the exact measurement conditions.

## Discussion

Our analysis confirmed our hypothesis that there is a set of sequence changes in the β-cardiac myosin, among mammals, that have a high probability of association with species mass. In the 67 species examined 52 sites in the motor domain the residue present varied in more than six species. Of these 52 sites a set of 24 sequence changes had the strongest association with mass (p<0.05) and little association with clade (p>0.05). These sites were found throughout the motor domain but we noticed three clusters of four residues (Fig 4) within the group that would allow a cloning approach to test if these residues do play a role in adjusting the velocity of muscle contraction. Of these 12 residues, nine differ between the rat and human β -myosin. We have an expression system for the human myosin motor domain and therefore made a human/rat chimera by exchanging these nine residues.

The chimera displayed a two-fold weakening of ADP binding to actin.myosin due to a two-fold acceleration of the rate constant controlling ADP release from the complex. The two fold faster ADP release rate constant, since it is believed to limit contraction velocity, predicts a two-fold acceleration of the velocity of muscle contraction and a two-fold acceleration of the speed at which actin would move over a bed of myosin. The motility assay confirmed this prediction. The mutations could have caused a generalised loss of nucleotide binding to the protein but a control examining the ability of ATP to bind to actin.myosin and displace actin was indistinguishable among the three proteins (Fig 7B).

Thus our bioinformatics approach has successfully identified nine residues with a role in modulating the velocity of muscle contraction which have been selected over time to adjust the velocity to that required for the slower contraction in human vs rat hearts.

Before considering the sites in the motor domain of myosin where these changes occur we should consider the role of the remaining 40 sites. As shown in Fig 4C, 21 sites have no apparent association with clade or mass and therefore the functional significance of these residue changes remains undefined. Eight sites have a strong association with clade but no or only modest association with mass. A further four sites had a strong association with both clade and mass. Each of these four sites that have strong association with both mass and clade along with the four sites, which were not made in the chimera, with a strong association with mass may contribute to the changes in contraction velocity between mammals. Our approach here was to establish the principle of the effect rather than delineate the contribution of every sequence change. Thus we focussed on three groups of amino acids

As stated in the introduction making single point mutations in the motor domain is unlikely to give sufficient experimental resolution to define the contribution of each residue to the 2-3 fold changes we were expecting between rat and human. However, groups of changes could possibly define the relative contributions of each of the three groups of residues. At the moment the complexity and expense of expressing the protein in mouse cell lines prohibits a larger scale study.

What was also unexpected was the finding that a set of ∼ 20 residues with the highest correlation with mass are predicted to have a narrow mass range over which each sequence change is found (Fig 6). Additionally, each of the residues has its own distinct midpoint for the transition. This implies that as mass increases, there is a limited number of ways in which the ADP release can be modified step-wise and mammals from distinct clades utilise the same set and order of sequence changes. This is a prime example of convergent evolution in the two clades.

None of the mutated residues are in direct contact with the nucleotide binding pocket (Fig 5), thus suggesting that the mutations have allosteric effects. Four of the changes occur in helix-O, a long helix in the upper 50 kDa domain that links the actin binding site (cardiomyopathy loop) with switch 2 in the nucleotide binding site. It is therefore in a position to influence the communication between these two important functional sites. However, the available crystal structures published from a variety of myosins show this helix to move with the whole of the upper 50 kDa domain (e.g. see Fig 7 in M. Bloemink et al., 2014). The other two groups or changes (three residues in the upper 50 kDa domain and two in the lower 50 kDa domain) are not close to each other in space. Understanding how the changes (for the most part conservative substitutions) in these regions alter the behaviour of the myosin will require a detailed molecular dynamics study. Seven of the twelve positions include amino acid changes that introduce or remove side chains that are capable of forming hydrogen bonds, thus it is possible that the sequence changes result in minor changes to hydrogen bonding networks in the protein. None of these sites are appear in the ClinVar web site as sites of mutations in human β-myosin associated with cardiomyopathies.

The results presented here show how a direct link between an organism’s physiology and a specific protein sequence allows the exploration of how selection may have adapted protein function to match the physiological requirements. The observation that the same set of amino acids have independently changed in two clades suggests constraints on the way a protein sequence can adapt whilst maintaining function. A wider study of muscle myosin sequences may show if different isoforms use the same or distinct sets of amino acids to adapt to the same selective pressure.

## Supporting information

Supplementary video 1 - In vitro motility of beta-cardiac and chimera S1. Scale bar is 5 microns

## Acknowledgements

NIH GM29090 to LAL; NIH HL117138 to LAL. MNW was supported by a Royal Society research grant. JM was supported by an EPSRC PhD studentship.

## Author Contributions

MAG, MNW, LAL and MR devised the study and supervised the research. JEM performed computational research. CAJ and DM planned the cloning strategy of the chimera. CAJ and CDV produced the chimera protein. CAJ performed biochemical characterisation of the proteins. CAJ and DM performed the motility assays. All authors contributed to the writing of the paper.

## Data availability

Source data files are available from Figshare - 10.6084/m9.figshare.8275406

## References

1. Bottinelli, R. & Reggiani, C. Human skeletal muscle fibres: molecular and functional diversity. Prog. Biophys. Mol. Biol. 73, 195–262 (2000).

2. Schiaffino, S. & Reggiani, C. Fiber Types in Mammalian Skeletal Muscles. Physiol. Rev. 91, 1447–1531 (2011).

3. Pellegrino, M. A. et al. Orthologous myosin isoforms and scaling of shortening velocity with body size in mouse, rat, rabbit and human muscles. J. Physiol. 546, 677–89 (2003).

4. He, Z. H., Bottinelli, R., Pellegrino, M. A., Ferenczi, M. A. & Reggiani, C. ATP consumption and efficiency of human single muscle fibers with different myosin isoform composition. Biophys. J. 79, 945–61 (2000).

5. Bloemink, M. J., Deacon, J. C., Resnicow, D. I., Leinwand, L. A. & Geeves, M. A. The superfast human extraocular myosin is kinetically distinct from the fast skeletal IIa, IIb, and IId isoforms. J. Biol. Chem. 288, 27469–79 (2013).

6. Resnicow, D. I., Deacon, J. C., Warrick, H. M., Spudich, J. A. & Leinwand, L. A. Functional diversity among a family of human skeletal muscle myosin motors. Proc. Natl. Acad. Sci. U. S. A. 107, 1053–8 (2010).

7. Savage, V. M. et al. Scaling of number, size, and metabolic rate of cells with body size in mammals. Proc. Natl. Acad. Sci. 104, 4718–4723 (2007).

8. Mcgreig, J. E. et al. Adaptation of mammalian myosin II sequences to body mass. 1–25 (2019). doi:https://doi.org/10.1101/055434

9. Walklate, J., Ujfalusi, Z. & Geeves, M. A. Myosin isoforms and the mechanochemical cross-bridge cycle. J. Exp. Biol. 219, 168–174 (2016).

10. Siemankowski, R. F., Wiseman, M. O. & White, H. ADP dissociation from actomyosin subfragment 1 is sufficiently slow to limit the unloaded shortening velocity in vertebrate muscle. Proc. Natl. Acadamy Sci. United States Am. 82, 658–662 (1985).

11. Homsher E, Wang F S. J. Factors affecting movement of F-actin filaments propelled by skeletal muscle heavy meromyosin. Am. J. Physiol. 263, 741–23 (1992).

12. Sievers, F. et al. Fast, scalable generation of high-quality protein multiple sequence alignments using Clustal Omega. Mol. Syst. Biol. 7, (2011).

13. R Core team. R: A language and environment for statistical computing. R Foundation for Statistical Computing, Vienna, Austria. URL https://www.R-project.org/. R: A Language and Environment for Statistical Computing. R Foundation for Statistical Computing, Vienna, Austria. ISBN 3-900051-07-0, URL http://www.R-project.org/. (2018). doi:10.2788/95827

14. Deacon, J. C., Bloemink, M. J., Rezavandi, H., Geeves, M. A. & Leinwand, L. A. Erratum to: Identification of functional differences between recombinant human α and β cardiac myosin motors. Cell. Mol. Life Sci. 69, 4239–55 (2012).

15. Huang, Jin. Nagy, Stanislav. Koide, Akiko. Rock, Ronald. Koide, S. A peptide tag system for facile purification and single-molecule immobilization. Biochemistry 48, 11834–11836 (2009).

16. Aksel, T., ChoeYu, E., Sutton, S., Ruppel, K. M. & Spudich, J. A. Ensemble Force Changes that Result from Human Cardiac Myosin Mutations and a Small-Molecule Effector. Cell Rep. 11, 910–920 (2015).

17. Spudich, J. A. & Watt, S. The regulation of rabbit skeletal muscle contraction. I. Biochemical studies of the interaction of the tropomyosin-troponin complex with actin and the proteolytic fragments of myosin. J. Biol. Chem. 246, 4866–71 (1971).

18. Criddle, A. H., Geeves, M. A. & Jeffries, T. The use of actin labelled with N-(1- pyrenyl)iodoacetamide to study the interaction of actin with myosin subfragments and troponin/tropomyosin. Biochem. J. 232, 343–9 (1985).

19. Margossian, SS. Lowey, S. Preparation of myosin and its subfragments from rabbit skeletal muscle. Methods Enzymol. 85, 55–71 (1982).

20. Walklate, J., Vera, C., Bloemink, M. J., Geeves, M. A. & Leinwand, L. The most prevalent freeman-sheldon syndrome mutations in the embryonic myosin motor share functional defects. J. Biol. Chem. 291, (2016).

21. Adhikari, A. S. et al. Early-Onset Hypertrophic Cardiomyopathy Mutations Significantly Increase the Velocity, Force, and Actin-Activated ATPase Activity of Human β-Cardiac Myosin. Cell Rep. 17, 2857–2864 (2016).

22. Johnson, M., East, D. A. & Mulvihill, D. P. Formins determine the functional properties of actin filaments in yeast. Curr. Biol. 24, 1525–1530 (2014).

23. Spudich, J. How molecular motors work. Nature 372, 515–518 (1994).

24. Bernstein, S. I. & Milligan, R. A. Fine Tuning a Molecular Motor : The Location of Alternative Domains in the Drosophila Myosin Head. J. Mol. Biol. 271, 1–6 (1997).

25. Miller, B., Bloemink, M. J., Geeves, M., Nyitrai, M. & Bernstein, S. A Variable Domain Near the ATP Binding Site in Drosophila Muscle Myosin is Part of the Communication Pathway between the Nucleotide and Actin-Binding Sites. J. Mol. Biol. 368, 1051–1066 (2007).

26. Qun Wang Carole L. Moncman and Donald A. Winkelmann. Mutations in the motor domain modulate myosin activity and myofibril organization. J. Cell Sci. 116, 4227–4238 (2003).

27. Rajani Srikakulam and Donald A. Winkelmann. Chaperone-mediated folding and assembly of myosin in striated muscle. J. Cell Sci. 117, 641–652 (2004).

28. Bloemink, M. J., Adamek, N., Reggiani, C. & Geeves, M. A. Kinetic Analysis of the Slow Skeletal Myosin MHC-1 Isoform from Bovine Masseter Muscle. J. Mol. Biol. 373, 1184–1197 (2007).

29. Nag, S. et al. Contractility parameters of human β-cardiac myosin with the hypertrophic cardiomyopathy mutation R403Q show loss of motor function. Sci. Adv. 1, e1500511 (2015).

30. Nyitrai, M. & Geeves, M. Adenosine diphosphate and strain sensitivity in myosin motors. Philos. Trans. R. Soc. B Biol. Sci. 359, 1867–77 (2004).

31. Ujfalusi, Z. et al. Dilated cardiomyopathy myosin mutants have reduced force-generating capacity. J. Biol. Chem. 293, 9017–9029 (2018).

32. Bloemink, M. et al. The hypertrophic cardiomyopathy myosin mutation R453C alters ATP binding and hydrolysis of human cardiac β-myosin. J. Biol. Chem. 289, 5158–5167 (2014).

33. Pereira, J. S. A. et al. Kinetic Differences in Cardiac Myosins with Identical Loop 1 Sequences. J. Biol. Chem. 276, 4409–4415 (2001).

34. Toniolo, L. et al. Expression of eight distinct MHC isoforms in bovine striated muscles: evidence for MHC-2B presence only in extraocular muscles. J. Exp. Biol. 208, 4243–4253 (2005).

