## Supplementary files for "Cardiac contraction velocity has evolved to match heart rate with body size through variation in β-cardiac myosin sequence"

Running title: Adaptation of β-cardiac myosin

Key words: evolution, motility, muscle

Address for correspondence:

Prof M.A.Geeves
School of Biosciences,
University of Kent,
Canterbury
CT1 7NJ
UK


Dr M.N. Wass
School of Biosciences
University of Kent,
Canterbury
CT1 7NJ
UK


Prof L.A. Leinwand
BioFrontiers Institute and
Department of Molecular, Cellular
and Developmental Biology,
University of Colorado Boulder,
Boulder CO 80309 USA


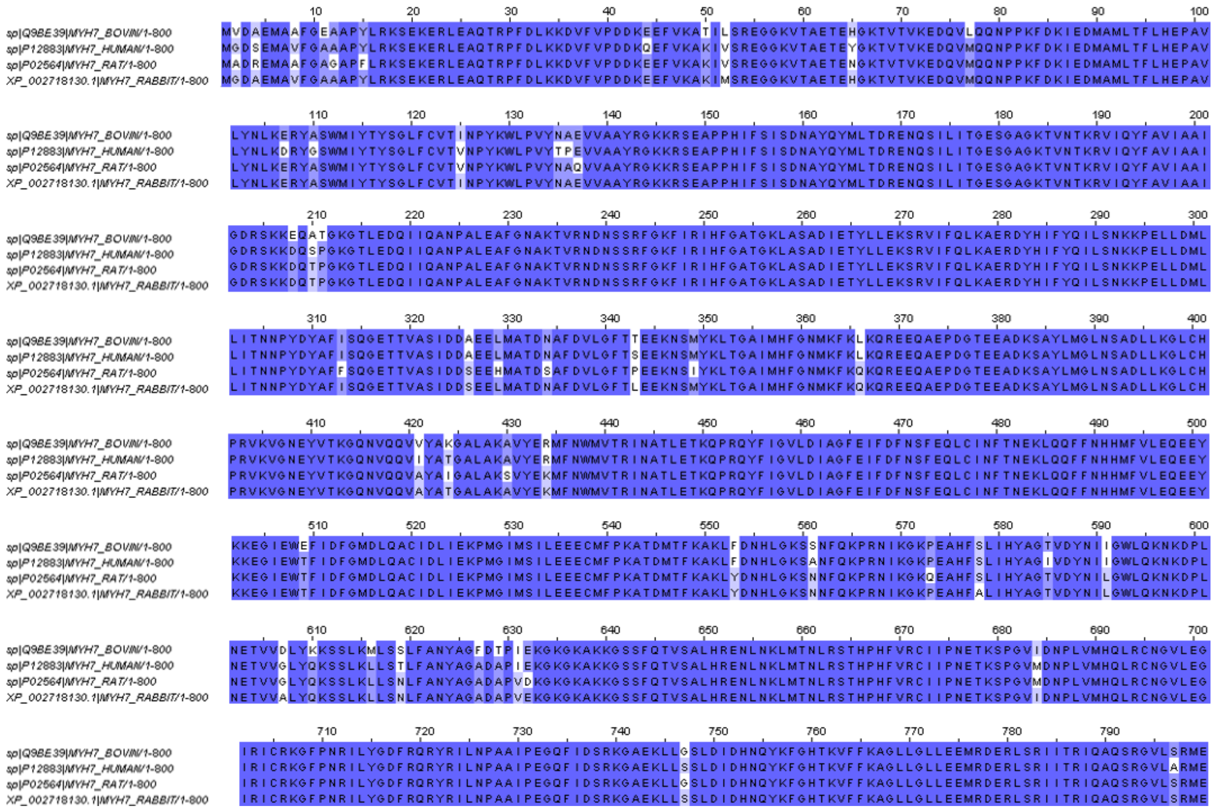

**Supplementary Figure 1. Multiple sequence alignment of rat, rabbit, human and bovine β-cardiac myosin.**

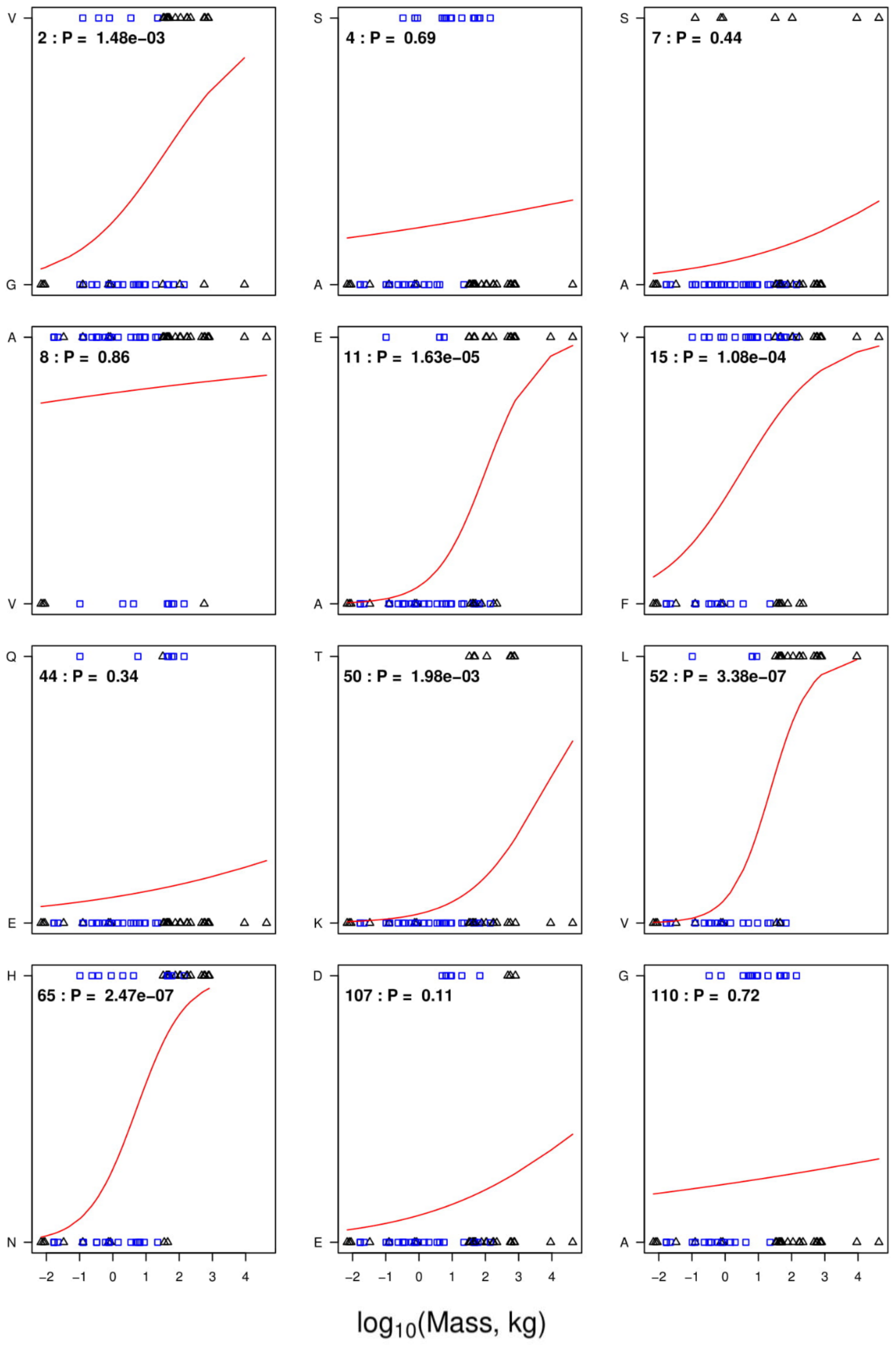
2 : P = 1.48e−03
4 : P = 0.69
7 : P = 0.44
8 : P = 0.86
11 : P = 1.63e−05
15 : P = 1.08e−04
44 : P = 0.34
50 : P = 1.98e−03
52 : P = 3.38e−07
65 : P = 2.47e−07
107 : P = 0.11
110 : P = 0.72
log10(Mass, kg)

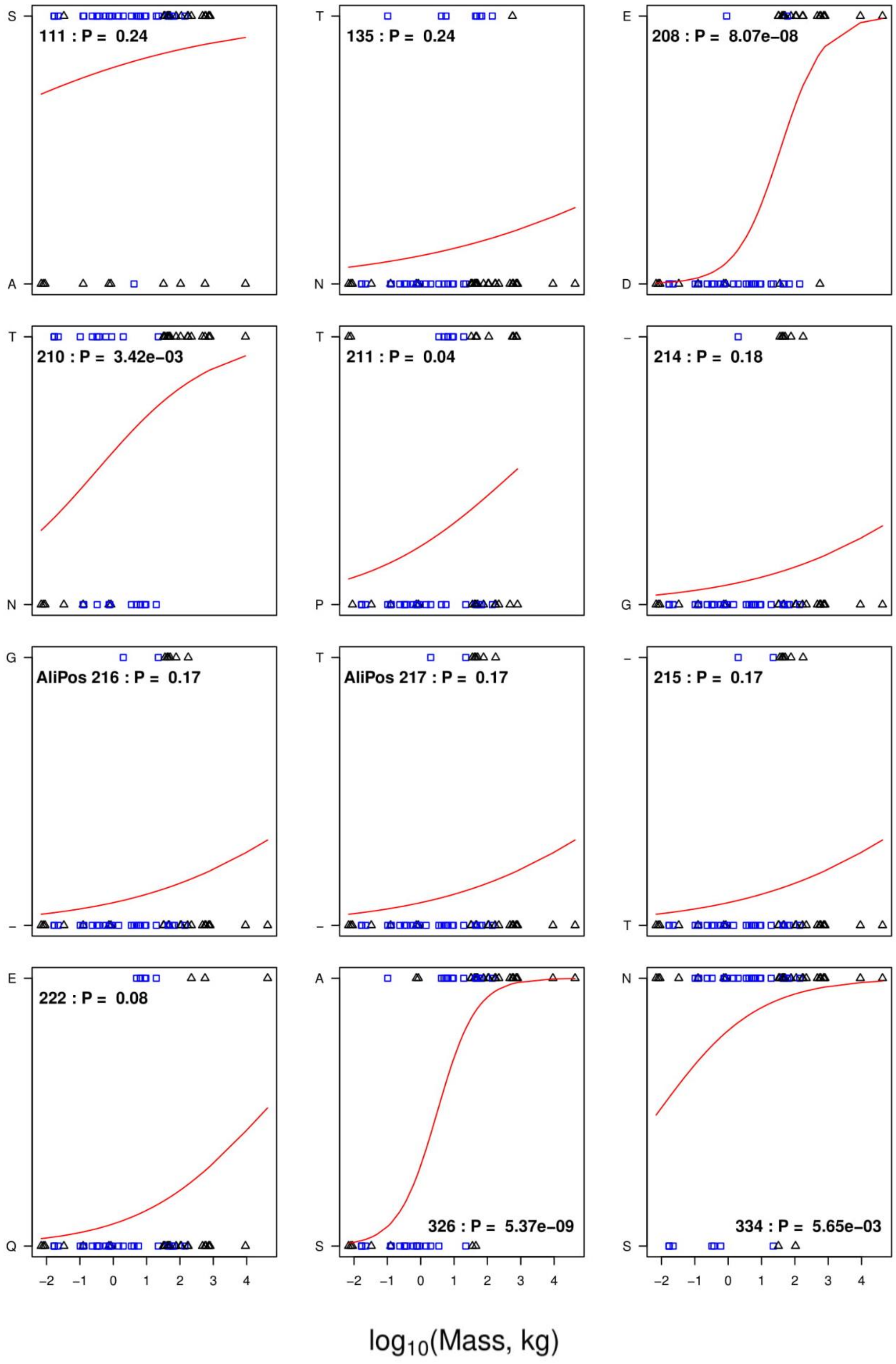

111 : P = 0.24
135 : P = 0.24
208 : P = 8.07e−08
210 : P = 3.42e−03
211 : P = 0.04
214 : P = 0.18
AliPos 216 : P = 0.17
AliPos 217 : P = 0.17
215 : P = 0.17
222 : P = 0.08
326 : P = 5.37e−09
334 : P = 5.65e−03
log10(Mass, kg)

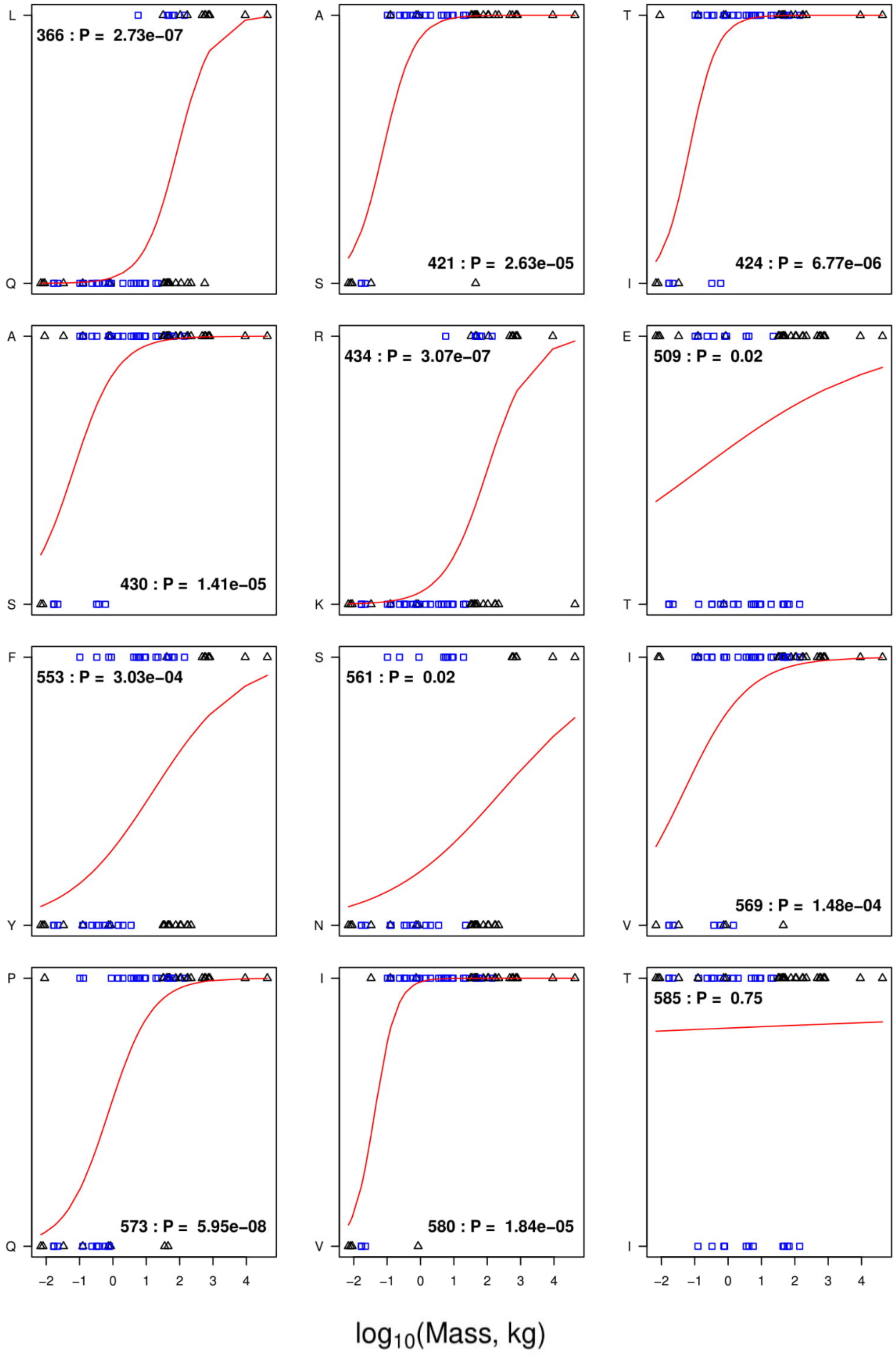

366 : P = 2.73e−07
421 : P = 2.63e−05
424 : P = 6.77e−06
430 : P = 1.41e−05
434 : P = 3.07e−07
509 : P = 0.02
553 : P = 3.03e−04
561 : P = 0.02
569 : P = 1.48e−04
573 : P = 5.95e−08
580 : P = 1.84e−05
585 : P = 0.75
L Q A S T I A S R K E T F Y S N I V P Q I V T I
−2 −1 0 1 2 3 4
log10(Mass, kg)

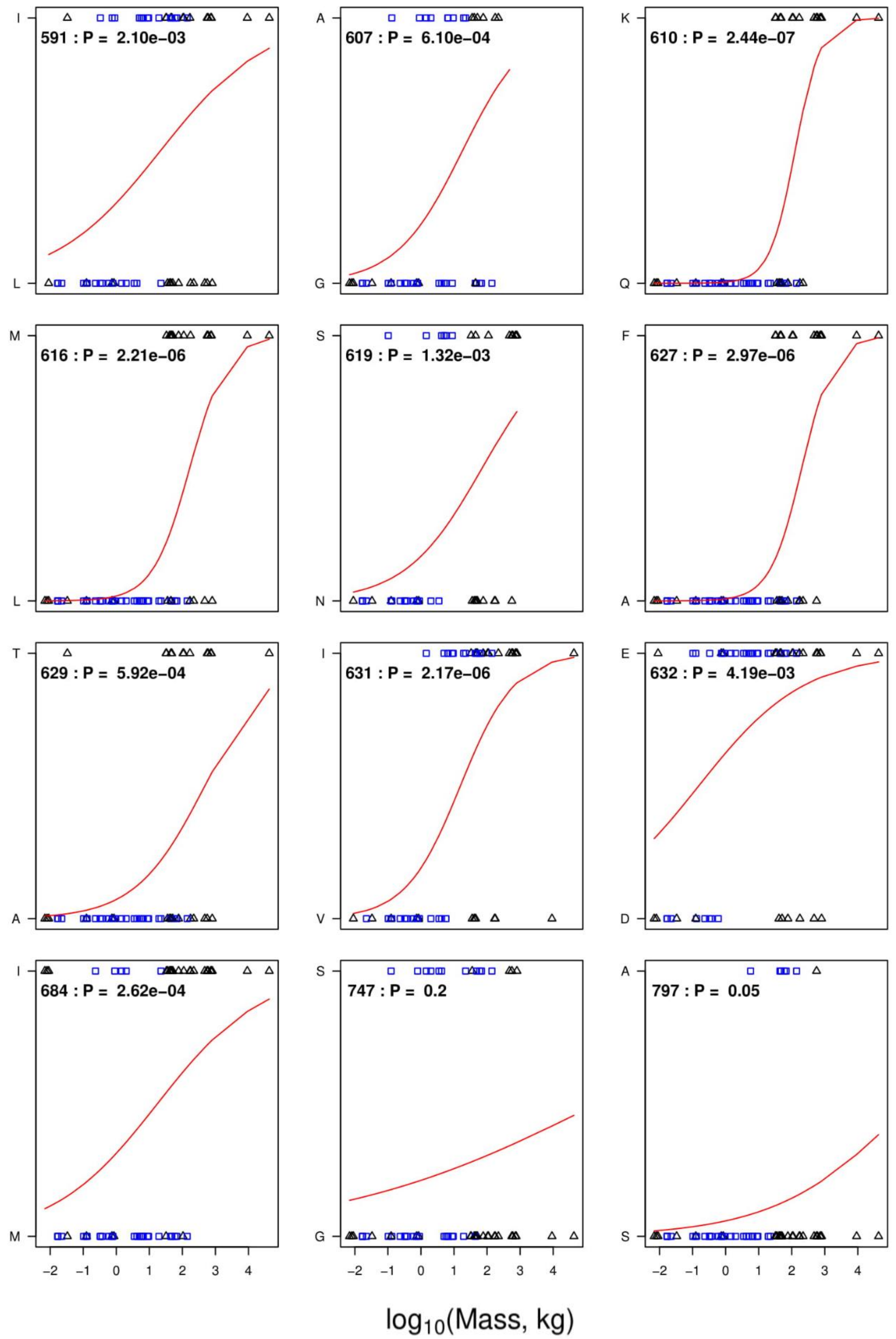


**Supplementary Figure 2. Residue-mass transition plots.** Binomial regression mapping the transition of the most frequent amino acid at positions in the motor region of β-myosin to the second most frequent amino acid at that position. The residue numbering is that of the human β-myosin, as oppose to the alignment position. The black squares are Euarchontoglires, and the triangles are Laurasiatheria. The P-value with each plot indicate the probability that the transition of the amino acids is not a result of change in mass.

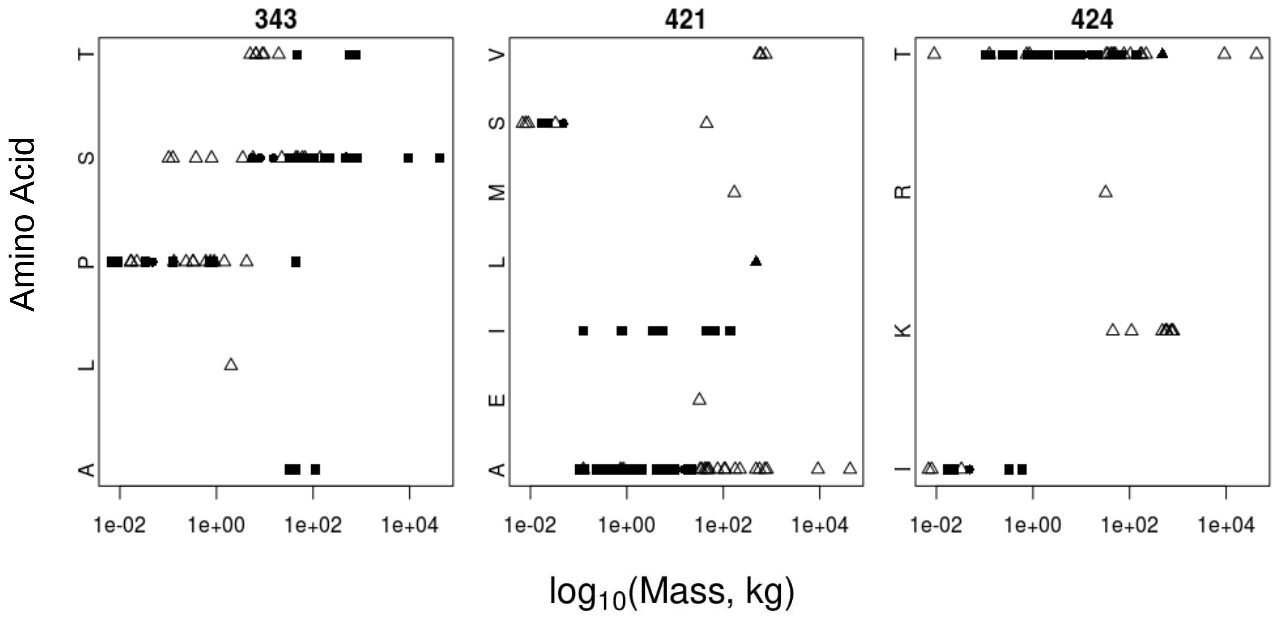


**Supplementary Figure 3. Highly variable residue mass vs amino acid frequencies.** Residues which had more than two sites of variation, with the third most frequent amino acid being close in frequency to the second most common amino acid. The black squares are Euarchontoglires, and the triangles are Laurasiatheria. The residue numbering is that of human β-cardiac myosin.

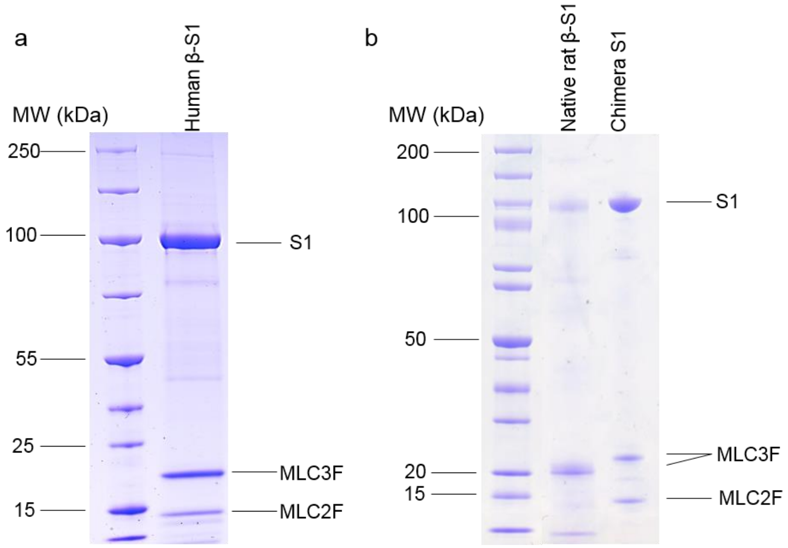


**Supplementary Figure 4. SDS-PAGE of the three protein preparations.** A. Recombinant human β-S1 with 2 light chains. B. Native rat β-S1 and recombinant chimera S1 with 2 light chains.

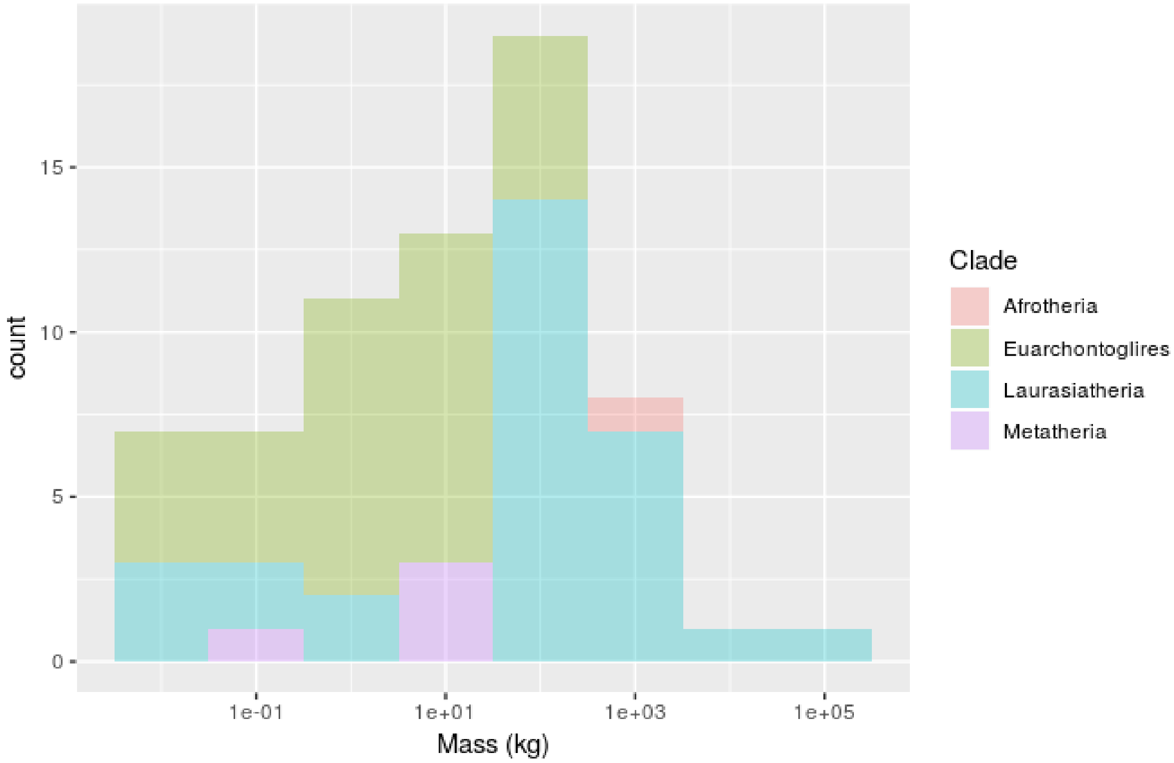


**Supplementary Figure 5. Frequency of organisms at different mass levels.** Plot to show the counts for organisms from each clade at different mass levels. Pink, dark, yellow, blue and purple shows organisms from the clades Afrotheria, Euarchontoglires, Laurasiatheria, and Metatheria clades.
